# Pharmacological Restoration of Visual Function in a Zebrafish Model of von-Hippel Lindau Disease

**DOI:** 10.1101/485730

**Authors:** Rebecca Ward, Kayleigh Slater, Zaheer Ali, Alison L Reynolds, Lasse D Jensen, Breandán N Kennedy

**Author notes:** **Correspondence:** Breandán Kennedy, UCD School of Biomolecular & Biomedical Science, UCD Conway Institute, University College Dublin, Belfield, Dublin D04 V1W8, Ireland.

## Abstract

Von Hippel-Lindau (VHL) syndrome is rare, autosomal dominant disorder, characterized by hypervascularised tumour formation in multiple organ systems. Vision loss associated with retinal capillary hemangioblastomas remains one of the earliest complications of VHL disease. The mortality of *Vhl*^*-/-*^ mice *in utero* restricted modelling of VHL disease in this mammalian model. Zebrafish harbouring a recessive germline mutation in the *vhl* gene represent a viable, alterative vertebrate model to investigate associated ocular *loss-of-function* phenotypes. Previous studies reported neovascularization of the brain, eye and trunk together with odema in the *vhl*^*-/-*^ zebrafish eye. In this study, we demonstrate *vhl*^*-/-*^ zebrafish almost entirely lack visual function. Furthermore, hyaloid vasculature networks in the *vhl*^*-/-*^ eye are improperly formed and this phenotype is concomitant with development of an ectopic intraretinal vasculature. Sunitinib, a multi tyrosine kinase inhibitor, market authorised for cancer, reversed the ocular behavioural and morphological phenotypes observed in *vhl*^*-/-*^ zebrafish. We conclude that the zebrafish *vhl* gene contributes to the endogenous molecular barrier that prevents development of intraretinal vasculature and that sunitinib can improve visual function and hyaloid vessel patterning while reducing abnormally formed ectopic intraretinal and choroidal vessels in *vhl*^*-/-*^ zebrafish.

## Introduction

Retinal hemangioblastomas, tumours of retinal blood vessels, are a characteristic clinical feature of Von Hippel Lindau (VHL) syndrome, a rare, systemic disease associated with significant loss of vision [1]. These benign, hypervascularised tumours represent the earliest complication of VHL disease, occurring in more than half of VHL patients. Development of retinal hemangioblastomas is dependent on dysregulation of ocular angiogenesis. Ocular angiogenesis is a dynamic process, controlled by an intricate balance of pro- and anti-angiogenic factors [2]. Imbalances can result in growth of abnormal, leaky vessels. VEGF isoforms and cognate receptors are key players which influence endothelial cell proliferation and migration during both developmental and pathological angiogenesis [3, 4].

Pathological neovascularisation is a significant feature of blinding disorders including diabetic retinopathy, retinopathy of prematurity and neovascular age-related macular degeneration [5]. Oncogenes, tumour suppressor genes and disruptions in growth factor activity promote pathological ocular angiogenesis. In VHL disease, loss-of-function germline mutations in the tumour suppressor gene *VHL* and subsequent inactivation of the second wildtype allele leads to aberrant hypoxia inducible factor-1 (HIF-1) expression and the development of multi-systemic tumours [1]. HIF-1 is a master transcriptional regulator, vital for cellular adaptation to hypoxia. In the absence of functional *VHL,* HIF-α translocates to the nucleus, upregulates transcription of pro-angiogenic mediators including VEGF and its receptors, erythropoietin (EPO), EPO receptor (EPOR), multidrug resistance pump (MDR-1) transferrin, angiopoietin 1, glucose transporters (GLUT-1) and glycolytic enzymes [6, 7].

Hypoxic pathways are well conserved between zebrafish and human making zebrafish a powerful model for hypoxia-related studies (Metelo et al., 2015). Furthermore, functional studies from forward and reverse genetics show the molecular components that regulate vascular development are conserved between mammals and fish [8] [9]. Two vascular networks form within the zebrafish eye during embryonic development – the hyaloid vessels located in the anterior eye, and the superficial choroidal vasculature which overlies the retinal pigmented epithelium and is responsible for oxygenation of the photoreceptors [10]. In contrast to zebrafish larvae, the hyaloid network in the human eye undergoes a programmed regression to form the retinal vasculature [11, 12].

A recessive *vhl* zebrafish model mimics some genetic and clinical hallmarks of VHL disease and provides a powerful model to study systemic and ocular-specific phenotypes linked to the absence of VHL signalling [13]. *vhl*^*-/-*^ zebrafish larvae display upregulated VEGF ligands (*vegfaa, vegfab* and *vegfd*) and receptors (*flt-1, kdr and kdr-like)* accompanied by ectopic cranial and intersegmental vasculature [14]. Aberrant expression of these HIF-target genes results in ectopic and leaky blood vessel formation in the trunk and eye, akin to pathological features observed in VHL patients [14]. Homozygous *vhl*^*-/-*^ zebrafish display pericardial and yolk oedema and typically die on or before day 13 due to cardiac or renal failure [13]. Previous pharmacological studies blocking HIF2α in *vhl*^*-/-*^ rescued embryonic pathologic angiogenesis and prolonged survival [7].

While the morphological vascular phenotypes of *vhl*^*-/-*^ zebrafish are characterised [7, 13-15] nothing is reported regarding the impact on visual function. Here, we identify and characterise an absent optokinetic response and a significantly reduced visual motor response in *vhl*^*-/-*^. We reaffirm the development of ectopic ocular blood vessels but clarify that this results from an attenuation of hyaloid vessels and the emergence of pathological ectopic intraretinal vessels in *vhl*^*-/-*^ larvae. We show a selective downregulation of v*egfaa* in *vhl*^*-/-*^ larval eyes following treatment with sunitinib malate, an anti-angiogenic compound approved for cancer treatment. Additionally, we demonstrate pharmacological improvement of visual function and normalisation of ocular vasculature in *vhl*^*-/-*^ following treatment. In summary, the *vhl* gene, by controlling *vegfaa* expression in the eye, prevents intraretinal vasculature development in zebrafish.

## Methods

### Zebrafish Breeding and Maintenance

All experiments carried out on animals were performed according to ethical approval granted by the UCD Animal Research Ethics Committee. Zebrafish were maintained in a 14-hour light, 10-hour dark cycle in a recirculating water system at 28^o^C and feed brine shrimp twice daily. *vhl*^*-/-*^ larvae were generated through natural spawning by incrossing *vhl*^*hu2117 +/-*^ zebrafish on transgenic background *Tg(fli1:EGFP)* and maintained as previously described [16].

### Genotyping

To identify adult *vhl*^*hu2117+/-*^ zebrafish, a section of the caudal fin was obtained under anaesthesia (Tricane MS-222) (Sigma Aldrich, UK). DNA was extracted using 50 mM NaOH (Sigma Aldrich, UK) before neutralising with Trizma Base (Sigma Aldrich, UK). The region of interest was amplified by PCR using the following primers which span the vhl mutation site: 5’-TAAGGGCTTAGCGCATGTTC-3’ and 5’-CTATCTACGCGTTTAACTCG-3’. PCR products were purified using a PCR clean up kit (New England Biolabs) and the single nucleotide polymorphism identified through sequence analysis (Source Biosciences, Ireland).

### Drug Preparation and Treatment

Sunitinib malate (Sigma Aldrich, UK) was added to embryo medium (0.137 M NaCl, 5.4 mM KCl, 5.5 mM Na2HPO4, 0.44 mM KH2PO4, 1.3 mM CaCl2, 1.0 mM MgSO4 and 4.2 mM NaHCO3) containing methylene blue. At 58 hpf, embryos were manually dechorionated and 400 µl of increasing test concentrations of sunitinib added to each well containing five embryos in a 48 well plate.

### Behavioural Analysis

To measure optokinetic response (OKR), larvae were immobilised in 9% methylcellulose (Sigma Aldrich, UK) and placed in the centre of a rotating drum containing black and white stripes. The drum was rotated at 18 rpm for 30 seconds in a clockwise and 30 seconds in a counter-clockwise direction. Saccades per minute were recorded manually (Deeti et al., 2014). To measure visual motor response (VMR), individual larvae were added to a square, flat bottomed 96 well plate in 600 µl of embryo medium and placed in a Zebrabox chamber (Viewpoint Life Sciences, France) to record locomotor activity in response to changes in light intensity. Larvae were acclimatised for 1 hour in light followed by 100 minutes of light in 20 minute ON-OFF intervals. Larval movement was quantified using Viewpoint software and data was analysed using a MS Excel spreadsheet as previously described [17].

### Hyaloid Vessel Analysis

5 dpf larvae were fixed in 4% paraformaldehyde overnight at 4°C and subsequently washed in 1X PBS-0.1% Tween 20 (PBST). Fixed larvae were screened for overall morphological defects before dissecting the lens. The number of eGFP positive primary HV and vessel branches on the lens posterior were quantified using an Olympus SZX16 fluorescent microscope.

### Intraretinal blood vessel analysis

Larvae were fixed in 4% PFA overnight at 4^o^C. Following 3 PBST washes, samples were transferred to a 5%:20% sucrose gradient before incubating overnight at 4^o^C in 20% sucrose. Samples were washed in 20% sucrose:OCT gradient before OCT embedding and storage at - 80^o^C. Samples cut to 20 µm sections were counterstained with DAPI. Images were taken using a Zeiss LSM510 confocal microscope (Carl Zeiss AG, Germany) with a 63X objective. Images were deconvoluted using Autoquant X3 software (Media Cybermetrics, USA) and projections generated using Imaris software (Bitplane, Switzerland). To quantify retinal vasculature, the area of eGFP+ blood vessels (minus the normal hyaloid artery) was manually measured using ImageJ software in sibling and *vhl*^-/-^ samples.

### Choriocapillaris development assay

Eggs were collected from Tg(*fli1a:EGFP*)^y1^ transgenic zebrafish and fertilization was confirmed at 2-4 hpf. Fertilized eggs were incubated in PTU-containing E3-medium for 58 or 72 hours as indicated and then treated with 0.1% DMSO (vehicle control) or 1 µM sunitinib malate (LC Laboratories) until 120 hpf. The larvae were then euthanized in 0.04% MS-222 (Ethyl 3-aminobenzoate methane sulfonic acid salt 98%, Sigma Aldrich) and fixed in 4% PFA (Sigma Aldrich) for 30 minutes at room temperature. The larvae were enucleated and the eyes dissected using watchmakers forceps (Dumont #5) under a dissection microscope (Nikon SMZ 1500). The retinae and choroids were flat mounted on glass slides in Vectashield mounting medium (H-1000 Vector laboratories) and imaged by confocal microscopy (Zeiss, LSM 700). The number of interstitial pillars (ISPs) were counted manually and used as a measure of the extent of active vascular growth.

### Histological Analysis

Larvae were fixed in glass vials containing 2.5% glutaraldehyde, 2% paraformaldehyde and 0.1% Sorenson’s phosphate buffer (pH 7.3) and placed at 4^o^C overnight. Samples were transferred to 1% osmium tetroxide before an ethanol gradient dehydration. Larvae were embedded in agar epoxy resin and sectioned using a glass knife and a Leica EM UC6 microtome. Sections were placed on glass slides and stained with Toluidine blue (Sigma Aldrich, UK) and imaged using a Leica DMLB bright field illumination microscope with a Leica DFC 480 camera.

### Total RNA extraction, cDNA synthesis and qPCR

Approximately 40 pooled larvae were stored in RNAlater (Qiagen) at 5 dpf before the eyes were enucleated and total RNA extracted using mirVana^TM^ miRNA Isolation Kit (ThermoFisher Scientific) as per the manufacturer’s instructions. Total RNA concentration was quantified at 260 nm (Spectrophotometer ND 2000) and samples stored at −80°C until further use. cDNA was synthesised using SuperScript II Reverse Transcriptase system (Invitrogen) using random hexamers as per supplier’s instructions.

RT PCR reactions were assembled on ice per manufacturer’s instruction using Phusion® High-Fidelity DNA Polymerase. (NEB). b-actin - 5’-CTTCCTGGGTATGGAATCTTGC-3’ and 5’-GTGGAAGGAGCAAGAGAGGTG-3’; vhl 5’TAGCTCAGAGACCCAGCATT-3’ and 5’-CAGTACATATGCACAGTCCAG-3’; sema3aa 5’-CTAAACAACAACAGCTGTACCTCG-3’ and 5’-CCTGACGTCTCGTTCTCCTT-3’; sema3ab 5’-GGATCTGCCATCGGGGTTT-3’ and TGACGCCTCGTTCTCCTCTTAG-3’; sema3fa 5’GGCACAGGGTTTTCTGCAAG-3’ and 5’-CCAGGCTCCAGTCGGAAAAT-3’; sema3fb 5’-TCTTCAAAACGGCAACAACTG-3’ and 5’- ACGGCTGCGTCTAAAAGGAA-3’.

To quantify *vegfaa, vegfab, vegfc* and *kdrl* expression, 0.5 μl of Taqman real time assay (Applied Biosystems) was used in each reaction.

### Statistical Analysis

Statistical analysis was performed using GraphPad Prism^TM^ software (GraphPad, San Diego, CA). The one-tailed student’s paired t-test was used when statistical analysis was required between two experimental groups. For analysis involving more than two independent groups with multiple test doses, a repeated measures one-way ANOVA and Dunnett’s multiple comparisons post hoc test was performed. When two variables were compared, a two-way ANOVA was used with Bonferroni post-test. All data are presented as mean ± standard error of the mean (SEM). Statistical significance was assigned to p-values of p < 0.05 = *, p < 0.01 = ** or p < 0.001 = ***.

## Results

### Characterisation of *vhl*^*-/-*^ zebrafish

Larvae were examined for key *vhl*^*-/-*^ phenotypes previously described [13]. Putative *vhl*^*-/-*^ larvae fail to inflate their swim bladder and display severe yolk and pericardial oedema at both 4 and 5 dpf (Figure 1.A). At 5 dpf, larvae from *vhl*^*+/-*^ incrosses were separated into two groups, predicted unaffected siblings and predicted *vhl*^*-/-*^ based on these phenotypes. Sequencing of individual larvae confirmed a complete correlation between phenotype and genotype. 100% of sibling larvae genotyped as *vhl*^+/+^ or ^+/-^ possessed an inflated swim bladder whereas this feature was absent in 100% of larvae genotyped *vhl*^*-/-*^ (Figure 1.B). The uninflated swim bladder phenotype method was used for rapid live selection of *vhl*^*-/-*^ larvae in all subsequent experiments. There was no significant difference in *vhl* mRNA expression in *vhl*^*-/-*^ larval eyes compared to sibling controls, indicating the absence of nonsense-mediated decay (Figure 1.E).

**Figure 1:**
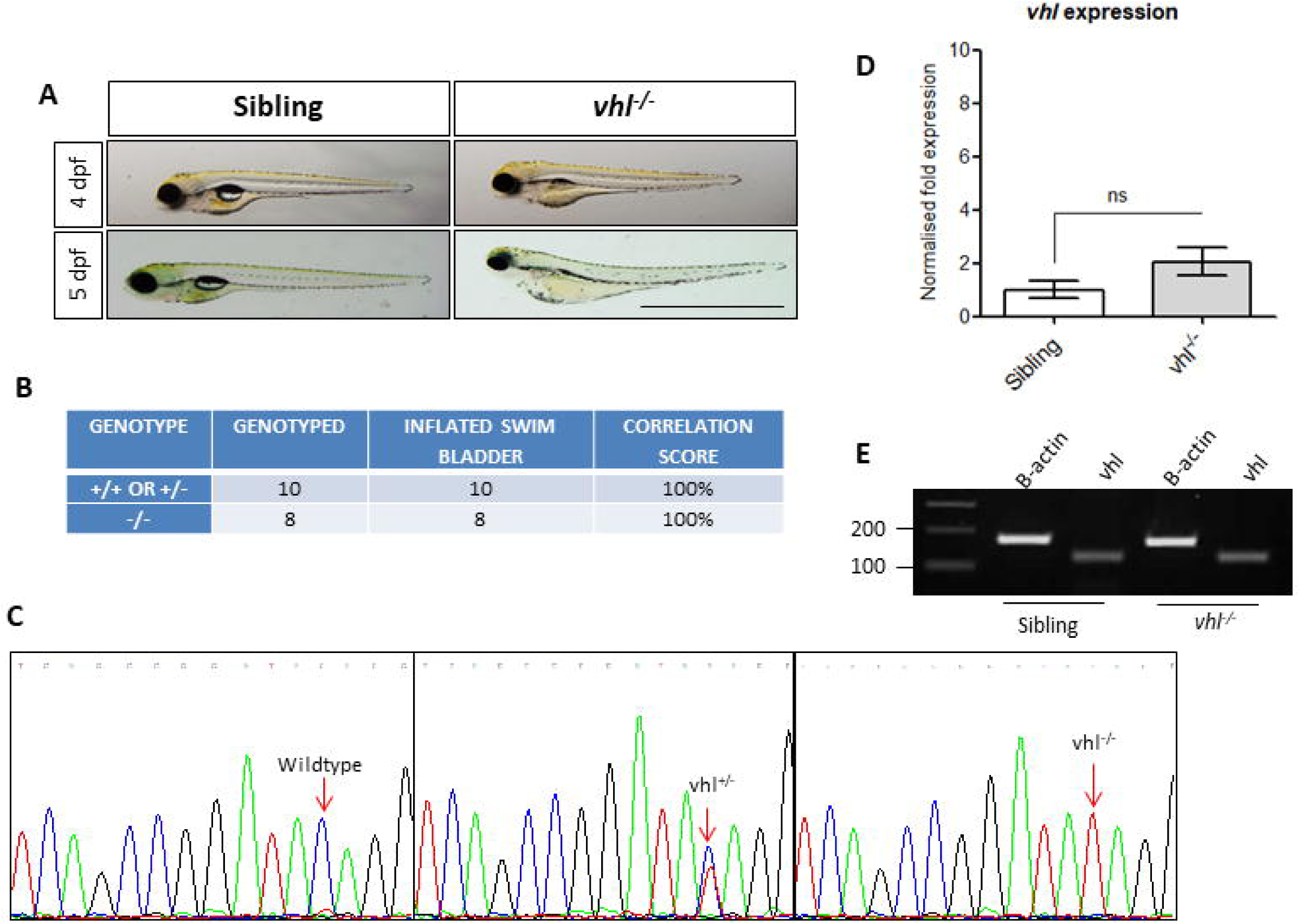
Characterisation of *vh*^*-/-*^ zebrafish. **A)** Gross morphology of 4 and 5 dpf larvae highlighting the absence of a swim bladder, plus yolk and pericaridial oedema in *vh*^*-/-*^ larvae. Scale bar = 2 mm **B)** Phenotype-Genotype correlations in vhl DNA sequencing of PCR amplicons from genomic DNA confirmed a positive correlation between swim bladder uninflation and *vh*^*-/-*^ genotype. This strategy was used to select *vh/*^*-/-*^ larvae for all further experimentation. **C)** Sequencing chromatograms illustrating the single nucleotide polymorphism (C>T) in the *vhl* sequence resulting in a premarure STOP codon. D) Real time PCR analysis of *vhl* expression in *vh/*^*-/-*^ in comparison to sibling (+/- or +/+) larvae. **E)** Reverse transcriptase PCR highlighting *vhl* expression in *vhl*^*-/-*^ and sibling eyes at 5 dpf.

### *vhl*^*-/-*^ larvae display retinal and visual function defects

The eye is a metabolically demanding organ [18]. Defects in *vhl*^*-/-*^ ocular vasculature were hypothesised to result in impaired vision. Two behavioural assays were applied to assess vision in *vhl*^*-/-*^ larvae. The OKR measures larval saccadic eye movements in response to moving black and white stripes, whereas, the VMR measures acute larval locomotor activity in response to lighting changes [17]. In *vhl*^-/-^, visual function is significantly reduced (p< 0.001) at 4 and 5 dpf with 95% and 94% reductions in the number of eye saccades, respectively (Figure 2.B). Moreover, unlike siblings, *vhl*^*-/-*^ have significantly attenuated (5-fold decrease in overall activity) responses to acute light illumination or extinguishment. Previous reports identified retinal detachment and neovascularisation in *vhl*^*-/-*^ retinal layers at 5.75 and 7.5 dpf [14]. To assess the onset of damage, *vhl*^*-/-*^ retinal sections were assessed at earlier time points. At 4 dpf, *vhl*^*-/-*^ appear to contain fluid around the lens compared to the unaffected sibling controls (Figure 2.Di-ii) however, there are no statistically significant differences between the thickness of retinal layers. At 5 dpf, *vhl*^*-/-*^ possess fluid accumulation throughout the retina and retinal detachment (Figure 2.D). Interestingly, morphometric analysis highlighted statistically significant differences in the ganglion cell layer and the retinal pigmented epithelium in 5 dpf *vhl*^*-/-*^ larvae.

**Figure 2:**
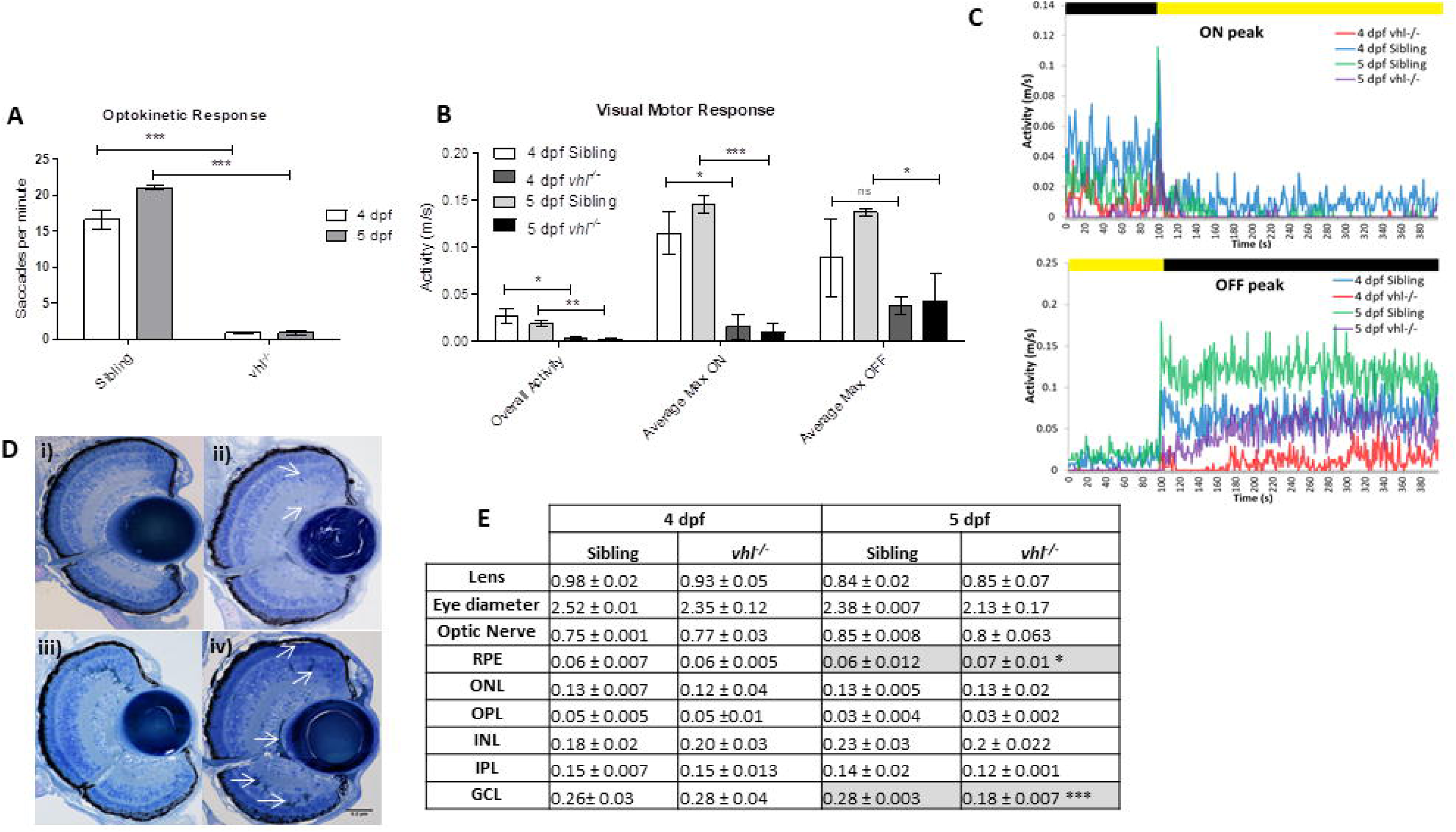
*vhl*^*-/-*^ larvae display retinal abnormalities and impaired visual function. *vhl*^*-/-*^ larvae possess **A)** reduced visual capacity in response to a moving striped stimulus (OKR) n=3, 12 technical replicates per treatment group per experiment. **B)** reduced movement in response to changes in light intensity (VMR) - VMR chromatographs illustrating average max ON/ OFF peaks. n=3, 12 technical replicates per treatment group per experiment. **D)** *vh*^*-/-*^ larvae display defects in retinal arrangementat 5 dpf. Representative light microscopy images of retinal sections from **i)** 4 dpf sibling control **ii)** 4 dpf *vhl*^*-/-*^ **iii)** 5 dpf sibling control and **iv)** extensive retinal damage, thought to be fluid accumulation present in 5 dpf *vhl*^*-/-*^ larvae (white arrows). **G)** Morphometric analysis based on 3 independent measurements from 3 larvae per group and shown as mean ± standard deviation. No statistical significant differences is observed between retinal layers of sibling and *vhl*^*-/-*^ at 4 dpf.

### *vhl*^*-/-*^ larvae possess retinal neovascularisation and lack hyaloid vessels at 4 and 5 dpf

To facilitate vasculature imaging, *vhl*^*hu2117*^ zebrafish were crossed into the Tg(*fli1*:EGFP)^y1^ line. Characterisation of ocular vessels in whole larvae by imaging through the physiological lens previously identified ectopic ocular vasculature in *vhl*^*-/-*^ [14]. To further characterise these vascular defects, *ex vivo* hyaloid vessel assays were conducted, wherein the lens is dissected from the retina and the number of primary eGFP^+^ blood vessels originating from the central point quantified [19]. Surprisingly, untreated *vhl*^*-/-*^ larvae contained significantly reduced primary hyaloid branches at 4 dpf (2-fold, p< 0.01) and 5 dpf (10.6-fold, p<0.001) compared to unaffected sibling controls (Figure 3 A-B). To locate the ectopic ocular vasculature, retinal cryosections were imaged by confocal microscopy. A specific pattern of ectopic intraretinal vascularisation was noted in *vhl*^*-/-*^ larvae with blood vessels branching throughout the inner plexiform layer at 4 dpf with more extensive growth 24 hours later (Figure 3.C). GFP positive areas in the retina, excluding the normal hyaloid artery were quantified by ImageJ. Intraretinal vessel area was significantly increased in *vhl*^*-/-*^ larvae at both 4 (37-fold, p< 0.01) and 5 dpf (68.2 fold, p< 0.01).

**Figure 3:**
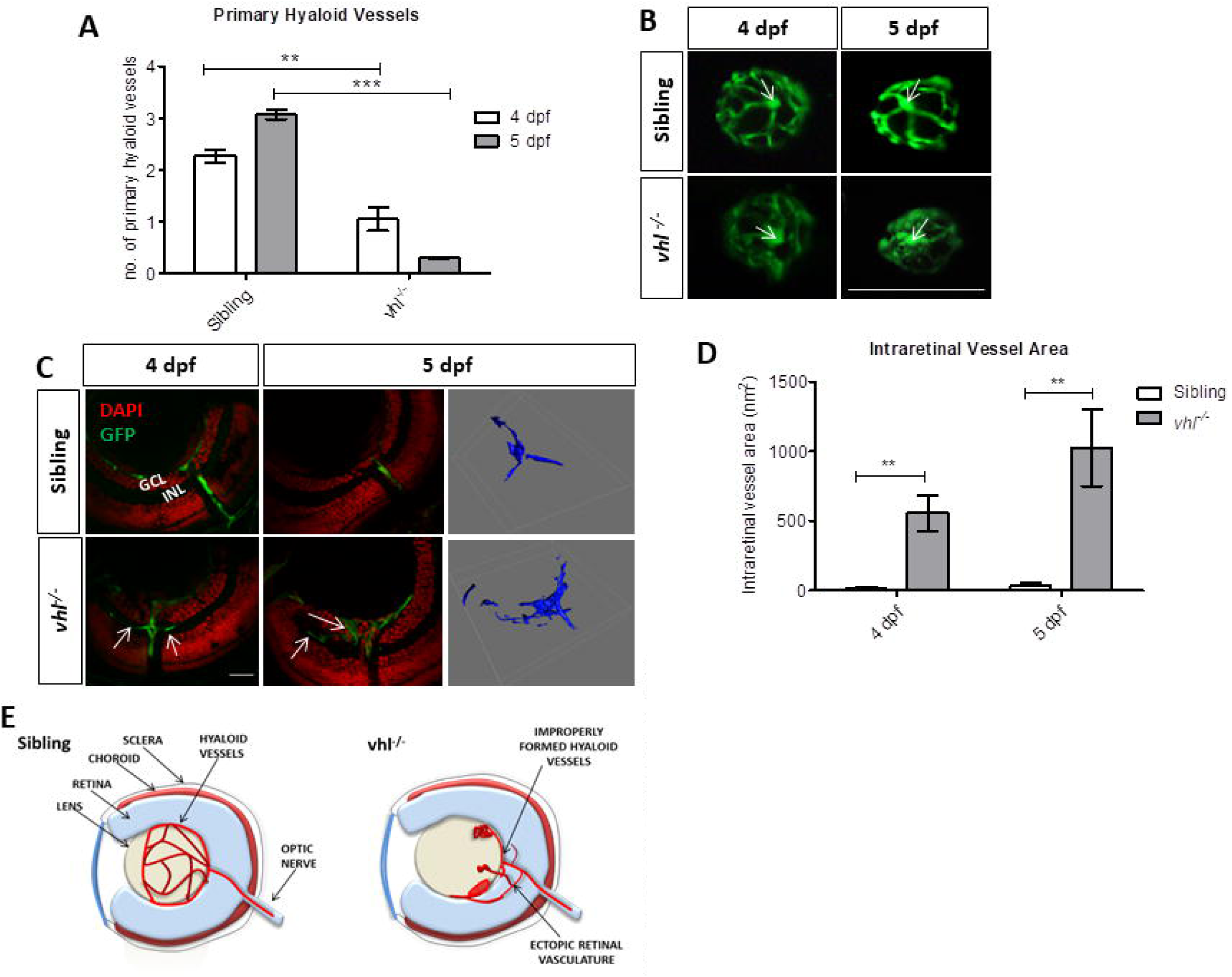
*vhl*^*-/-*^ zebrafish larvae possess vascular abnormalities at 4 and 5 dpf. **A)** Primary hyaloid vessel number at 4 and 5 dpf. *vhl*^*-/-*^ larvae highlight a significant reduction in hyaloid branches at 4 and 5 dpf compared to sibling control. Data is representative of 3 biological replicates. N=10 larvae per group per biological replicate. **B)** Fluorescent images of *fli1*:EGFP hyaloid vessels on sibling and *vhl*^*-/-*^ dissected lenses at 4 and 5 dpf. White arrows illustrate central point from which primary branches are quantified. Scale bar = 200 µm. **C)** Confocal imaging revealed abnormal retinal neovascularisation (white arrows) at 4 and 5 dpf through the inner plexiform layer (IPL) and ganglion cell layer (GCL) compared to sibling control. Z-stacks confirmed aberrant vessel growth (blue) through the retina. Scale bar = 20 µm. **D)** Intraretinal vessel area is significantly increased in *vhl*^*-/-*^ at both 4 and 5 dpf. N=5 larvae per group **E)** Schematic of proposed ocular blood vessel formation in *vhl*^*-/-*^ zebrafish larvae at 5 dpf.

### Sunitinib malate significantly improves visual function and reduces VEGF expression in *vhl*^*-/-*^ larvae

We hypothesised that pharmacological agents known to reduce angiogenesis, oedema, inflammation or neurodegeneration [14, 20, 21] may restore vision in *vhl*^*-/-*^ larvae. However, a histone deacetylase inhibitor (trichostatin A) and a corticosteroid (dexamethasone) failed to improve visual function or normalise hyaloid vessel development (*data not shown*). In contrast, treating *vhl*^-/-^ larvae with an anti-angiogenic (sunitinib) in a dose-response range of 0.2- 2.5 µM demonstrated that 1 µM sunitinib improved OKR visual behaviour at 5 dpf producing maximal improvement with a ∼20-fold increase (p<0.001) in visual function (Figure 4.B). *vhl*^*-/-*^ larvae treated with 1 µM sunitinib also resulted in improved VMR, but non-significantly, locomotor response to lights ON (3 fold, p>0.05) and OFF (1.4 fold, p> 0.05). Conversely, 0.5 μM sunitinib is capable of significantly increasing larval response to lights ON (p< 0.05) (Figure 4.D-E). Treating wildtype larvae with increasing doses of sunitinib (0.25 μM – 10 μM) reduced visual function by OKR and primary hyaloid vessel number in a dose dependent manner (Figure 4.C). This indicates that visual function is correlated to hyaloid vessel development.

**Figure 4:**
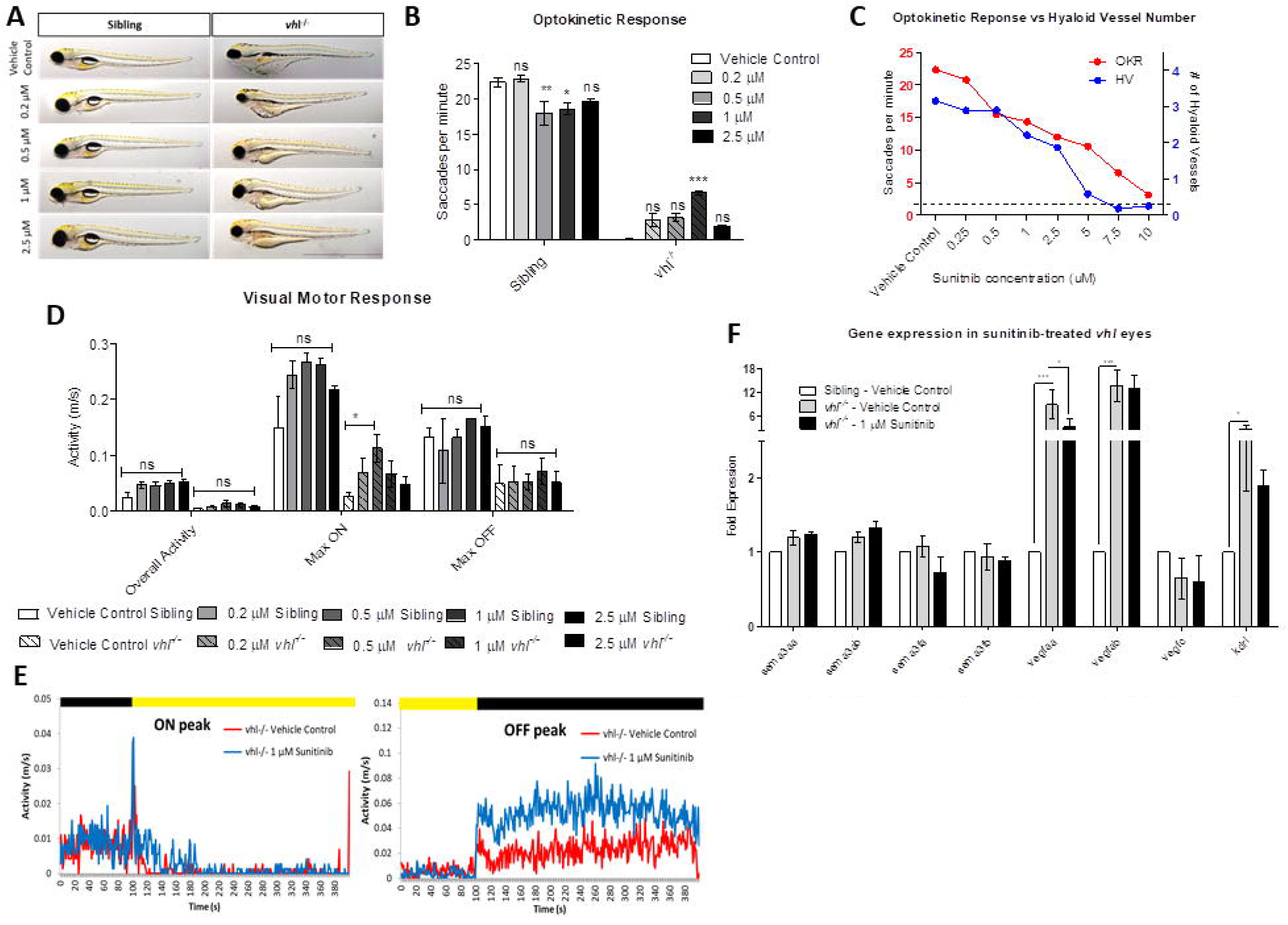
Sunitinib malate significantly improves visual function and reduces VEGF expression in *vhl*^*-/-*^ larvae. **A)** Gross morphology of larvae treated with increasing doses of sunitinib malate at 5 dpf. Treatment reduces yolk and cardiac oedema but does not result in swim bladder inflation. **B)** All concentrations of sunitinib tested increase OKR visual function at 5 dpf with 1 µM highlighting a statistically significant improvement. n=3, 12 technical replicates per experiment. **C)** Correlation between hyaloid vessel number and visual function in 5 dpf wildtype larvae. n=3 with 10 technical replicates per experiment. Dotted line denotes *vhl*^*-/-*^ visual function and munber of primary hyaloid branches at 5 dpf. **D)** Max ‘ON’ response is significantly increased dose dependently in larvae treated with 0.2 µM and 0.5 µM Sunitinib. n=3, 12

To elucidate molecular processes contributing to improved *vhl*^-/-^ vision, ocular mRNA levels of pivotal pro- and anti-angiogenic regulators were quantified. mRNA expression of all *sema3* isoforms in *vhl*^*-/-*^ eyes, before or after sunitinib treatment, were comparable to sibling control eyes. *vegfaa* expression is significantly upregulated in *vhl*^*-/-*^ fish, suggesting it is a driver of the *vhl*^*-/-*^ phenotype [14]. Although not restored to sibling levels, *vegfaa* expression was significantly reduced (5.6-fold, p< 0.05) following 1 μM sunitinib treatment (Figure 4.F). In contrast, the expression of *vegfab and kdrl* are significantly upregulated in the *vhl*^*-/-*^ eye however expression levels were unchanged by sunitinib treatment (Figure 4.F).

### Sunitinib malate improves hyaloid vessel patterning, reduces retinal neovascularisation normal and inhibits excessive development of the choriocapillaris in *vhl*^*-/-*^ larvae

We hypothesised that restored visual function and reduced ocular *vegfaa* levels following sunitinib treatment may correlate with improvements in the *vhl*^-/-^ vascular morphology phenotypes. Interestingly, the anti-angiogenic drug did not decrease primary hyaloid branch number. Additionally, while it did not significantly increase primary hyaloid branch number, sunitinib normalised patterning of the hyaloid vessels at all concentrations (0.2 – 2.5 µM) tested (Figure 5.B). In contrast, the area of ectopic intraretinal vascularisation was significantly reduced (3-fold, p< 0.05) following treatment with 1 μM sunitinib from 58 hpf (Figure 5.C). We, therefore, also investigated the integrity of the choroidal vessels. Compared to unaffected siblings, the number of interstitial pillars (ISPs) in *vhl*^*-/-*^ fish is significantly increased in number while decreased in size (1.6-fold, p< 0.05 and 4.4-fold, p< 0.01 for larvae treated at 72 and 58 hpf, respectively), leading to increased density of the choriocapillaris (Figure 5.D and E). There was no evidence of ectopic growth of vessels into the *vhl*^-/-^ neuroretina from the choriocapillaris. Treating larvae with sunitinib from 72 - 120 hpf, significantly reduced the number of ISPs in both wildtype and *vhl*^*-/*^ (Figure 5.D and E). Interestingly, starting treatment at 58 hpf led to a much more pronounced, anti-angiogenic effect in the choriocapillaris, particularly in the *vhl*^*-/-*^ larvae (Figure 5.D and E).

**Figure 5:**
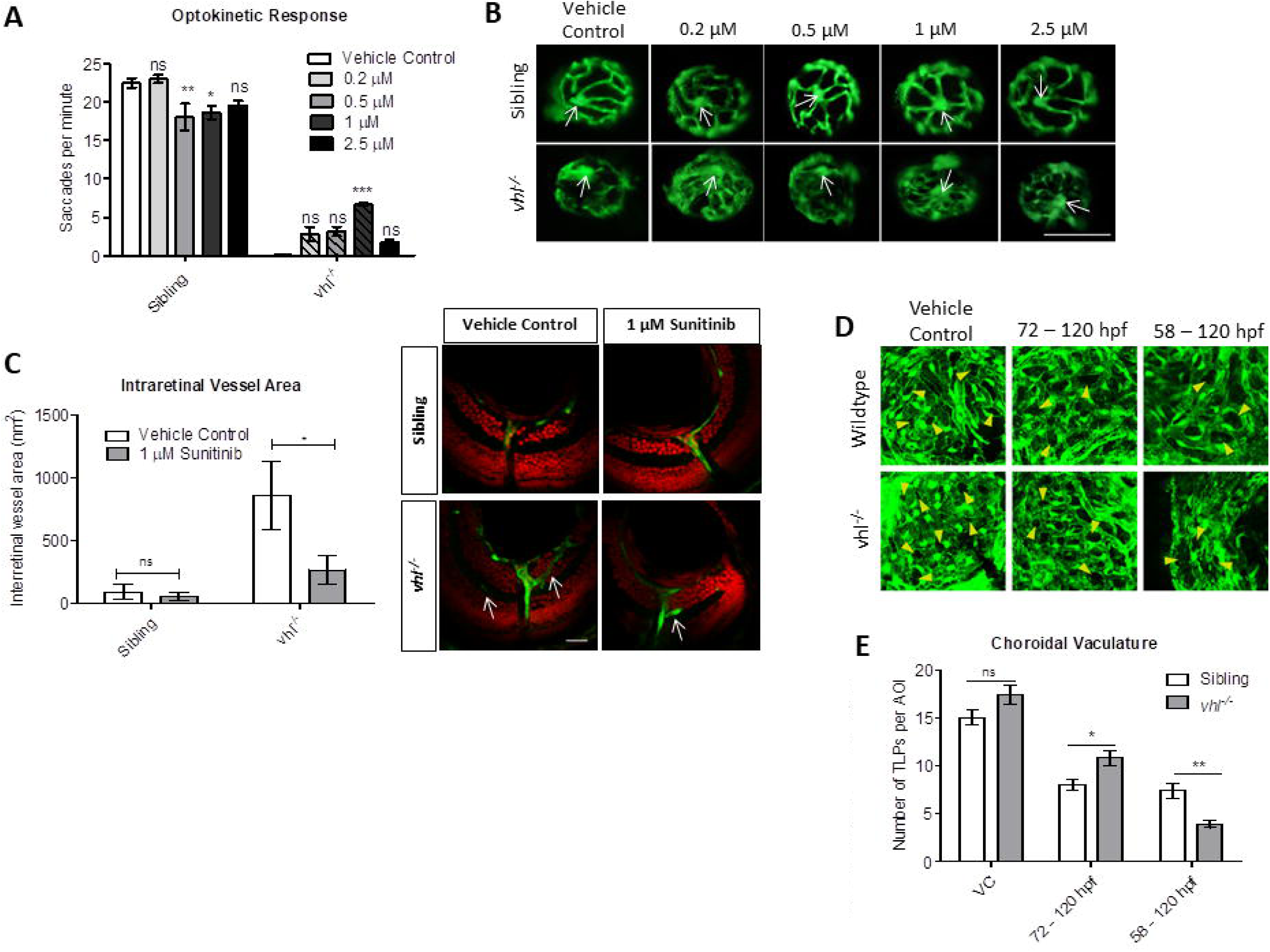
Sunitinib malate improves hyaloid vessel patterning, reduces retinal neovascularisation and inhibits excessive development of the choriocapillaris in *vhl*^*-/-*^ larvae. **A)** Treatment from 58 hpf does not significantly increase *vhl*^*-/-*^ primary hyaloid vessel nwnber at 5 dpf. **B)** Representative images highlighting an overall improvement of vessel patterning following treatment with Swiitinib from 58 hpf. White arrows denotes central point from which primary blood vessels are counted. n=3, 12 technical replicates per group per experiment. **C)** Reduction of ectopic intraretinal vasculature following treatment with 1 µM Sunitinib. White arrows highlighting ectopic intraretinal vessels. Quantification of n=5 larvae per treatment group. Scale bar = 20 µm. **D)** Z-stack projections of the sub-retinal choriocapillaris vasculature (endothelial cells shown in green) of Tg(*fli1*:EGFP) or *vhl*^*-/-*^ (*fli1*:EGFP) at 120 hpf following treatment for 72 or 48 hours as indicated. Yellow arrowheads point to interstitial pillars (ISPs). **E)** Quantification of the number of ISPs per area of interest (AOI). n=10 larvae.

## Discussion

Here, we report original findings on ocular phenotypes in the zebrafish *vhl*^*-/-*^ model. We demonstrate for the first time, profoundly attenuated visual behaviour in *vhl*^*-/-*^ larvae. In addition, we prove unequivocally that *vhl*^*-/-*^ possess extensive ectopic intraretinal vasculature and abnormal hyaloid vessel patterning. These phenotypes occur in parallel with elevated *vegfaa* levels in the eye. Significantly, treatment with the potent anti-angiogenic agent sunitinib culminated in reduced ocular *vegfaa* levels, resolved ocular vasculature dysmorphology and restored visual function.

Visual capacity in 4 and 5 dpf *vhl*^*-/-*^ larvae is significantly reduced when examined by OKR and VMR behavioural assays. We hypothesised that the aetiology of this visual defect was due to ocular vascular phenotypes. Retinal vasculature is absent at 5 dpf in zebrafish [22]. Instead, at this stage, the eye is nourished by the choroidal and hyaloid vascular networks [11]. Ectopic vessel growth in *vhl*^*-/-*^ zebrafish eyes was previously noted by imaging whole animals through the lens [14]. Here, dissections of larval lenses were performed to quantify *ex vivo* the number of primary hyaloid vessels positioned on the posterior lens. Surprisingly, the primary hyaloid vessel number was significantly reduced in *vhl*^*-/-*^ larvae. Subsequent analysis of cryosections revealed that the ectopic blood vessel formation was within the *vhl*^*-/-*^ retina. 3D confocal image projections show the ectopic vasculature to grow juxtaposed within the inner plexiform layer. This suggests an aberrant upregulation of pro-angiogenic factors or loss of an anti-angiogenic barrier in the *vhl*^*-/-*^ retina causing the vessels to turn and grow into the IPL rather than only projecting out to the lens. Interestingly, there were no signs of vessel growth from the choriocapillaris in the *vhl*^*-/-*^ neuroretina. This could potentially be explained by the presence of an extra-vascular barrier known as Bruch’s membrane separating the choriocapillaris from the neuroretina, which prevents the ingrowth of vessels into the outer retina also in humans [23].

It remains unclear as to exactly why *vhl*^*-/-*^ are visually impaired. The intraretinal vasculature could provide a physical barrier to light reaching the photoreceptor layer which appears structurally intact in histological sections. In addition to this, lack of properly formed or absent hyaloid vessels may reduce visual function as shown here in wildtype larvae when treated to higher doses of sunitinib malate. Although *vhl*^*-/-*^ fish are not hypoxic, their loss of vision may be adaptation to the systemic upregulation of hypoxic factors, similar to the crucian carp which is documented to reversibly lose its vision under anoxic conditions as an energy saving mechanism [24].

Difficulty modelling VHL disease in rodents due to embryonic lethality has stunted the progress towards elucidating the molecular mechanisms contributing to retinal hemangioblastoma growth and ultimate vision loss in humans. A conditional RPE-specific Vhl knockout mouse model developed aniridia, microphthalmia, retinal neovascularisation and RPE-specific cell death [25]. Additional deletion of HIF-α resulted in proper iris development and normal ocular growth, suggesting these processes require VHL-dependent regulation of HIF-α in the RPE. Moreover, *Vegf* expression and the development of abnormal retinal neovascularisation persisted in the double knockout mouse which suggests a HIF-independent regulation of vascular formation in the mouse retina [25]..

Previous pharmacological studies in *vhl*^*-/-*^ zebrafish with HIF2α inhibitors and anti-angiogenic compounds prolonged survival and reduced ectopic vasculature formation, respectively (Metelo et al., 2015; van Rooijen et al., 2010). A low dose of sunitinib was tested in line with previous findings demonstrating that 0.2 μM was sufficient to reduce ectopic vessel growth in *vhl*^*-/-*^ zebrafish trunk [14]. In this study, restoration of visual function and normalisation of hyaloid vessel patterning was observed with 0.2 μM sunitinib and increasing to 1 μM sunitinib produced the most significant rescue of vision in *vhl*^*-/-*^. Treating *vhl*^*-/-*^ with 1 μM sunitinib from 58-120 hpf, i) reduced the ectopic intraretinal vascular area, ii) reduced choroidal vessel interstitial pillars and iii) normalised patterning of the hyaloid vasculature. As the choriocapillaris is lumenized and perfused between 48 and 72 hpf, inhibiting tyrosine kinase activity during this developmental window may therefore more severely affect vascular survival and maturation. Additionally, the choriocapillaris was affected in wildtype indicating the requirement of this tyrosine kinase signalling in choroidal vasculature development. It is difficult to explain the differential effects of sunitinib treatment on different ocular vessel networks. It is possible that development of ectopic retinal vasculature results from aberrant *vegfaa* expression in the *vhl*^*-/-*^ retina. Sunitinib mediated *vegfaa* downregulation may be sufficient to reduce vessel growth in the retina and encourage subsequent hyaloid vessel development. It is likely that other effects of hypoxia signalling present in the *vhl*^*-/-*^ larvae (e.g. altered metabolism) may synergize with the lack of growth factor signalling in sunitinib-treated *vhl*^*-/-*^ larvae, leading to a more robust vascular disrupting effect in the intraretinal and choroidal vessels than the hyaloid network. In the retina, *VEGF* expression is highly controlled both spatially and temporally [26]. VEGF-A is secreted by the lens during mouse embryogenesis indicating a role in stimulating vessel growth. In *vhl*^*-/-*^, loss of *vegf* regulation may lead to the formation of intraretinal vessels and loss of hyaloid vessel development. Reducing *vegfa* expression, particularly *vegfaa* may be sufficient to reduce aberrant intraretinal growth and stimulate hyaloid vessel development.

Sema3A and Sema3F are expressed in the inner and outer mouse retina, respectively [27]. Due to their known vasorepressive properties in the retina, we evaluated expression of *sema3a (a/b)* and *sema3f (a/b)* isoforms in the *vhl*^*-/-*^ eye following treatment. No difference in *sema3* expression was documented before or after treatment with 1 μM sunitinib indicating other anti-angiogenic factors preventing the development of retinal vasculature may be lost in *vhl*^*-/-*^ zebrafish. As potent pro-angiogenic mediators, *vegf* expression was tested. A significant reduction was detected in *vegfaa* following treatment indicating that *vegfaa* is important in driving retinal angiogenesis in *vhl* zebrafish.

Development of specific and effective therapeutics for reducing ocular neovascularization depends on gaining a deeper understanding of the processes leading to vessel growth. VEGF is a well described pro-angiogenic factor involved in blood vessel formation through homing of circulating mononuclear myeloid cells and perivascular positioning and retention [28]. Clinical experiences with anti-VEGF agents for the treatment of wet-AMD have shown remarkably beneficial effects [29].

VHL disease is a highly-penetrant tumour predisposition syndrome in which multiple organ systems are affected. Development of retinal hemangioblastomas is a significant cause of visual morbidity in VHL patients. These lesions are normally associated with retinal neovascularisation through formation of arterial feeder vessels [30]. Currently there are no effective drug therapies for VHL disease and treatment of retinal hemangioblastomas relies on repeated surgeries or laser photocoagulation [30]. Our data demonstrates that reduction of ectopic ocular neovascularisation with sunitinib can restore vision in *vhl*^*-/-*^. However, direct translation to the clinic is inappropriate as *i)* the zebrafish knockout model does not recapitulate the development of retinal hemangioblastomas presenting in VHL patients and *ii)* resistance and systemic toxicity are major barriers to the clinical use of sunitinib [31]. Previous clinical trials evaluating the use of sunitinib to treat advanced retinal hemangioblastomas documented that treatment did not improve visual acuity or reduce the size of the lesion [32]. Nonetheless, the *vhl*^*-/-*^ zebrafish model provides a reliable, vertebrate model to study pathological angiogenesis and screen for genes and drug compounds that regulate intraretinal neovascularisation.

## Supporting information

## Acknowledgements

We thank UCD Conway Institute genomics facility for assistance with QRT PCR data analyses and UCD Conway Institute Imaging Core, with particular thanks to Dimitri Scholz and Tiina O’Neill. We also wish to thank Prof. Cormac Taylor for some helpful discussions surrounding this project and Elizabeth Young for technical assistance. This project was supported by Fighting Blindness MRCG/HRB grant MRCG-2014-3.

## Author Contributions

R.W., analysis of visual function, ocular vasculature, retinal histology and drug treatment experiments. R.W. and K.S., gene expression analysis. Z.A. and L.D.J., Analysis of choroidal vessels. R.W. and B.N.K., experimental design. R.W., A.L.R and B.N.K., Interpretation of results. R.W and B.N.K. wrote the paper. All authors approved the final manuscript.

